# A novel therapeutic approach of ultrasound stimulation to restore forelimb functions following cervical cord injury in rats

**DOI:** 10.1101/2022.10.03.510697

**Authors:** Rakib Uddin Ahmed, Monzurul Alam, Shuai Li, Poornima Palanisamy, Hui Zhong, Yong-Ping Zheng

## Abstract

Low-intensity pulsed ultrasound (LIPUS) stimulation has shown promising results in neurorehabilitation following a traumatic injury in brain and peripheral nerves. However, the effects of LIPUS stimulation in the injured neural circuit after spinal cord injury (SCI) are still unknown. We investigated the effects of LIPUS on forelimb functions in chronic cervical cord injured rats with and without a serotonergic agonist drug, Buspirone treatment. Twenty-six rats were trained for forelimb reaching and grasping followed by C4 dorsal funiculi crush injury. To deliver LIPUS, a silicon-coated ultrasound disc was implanted above the cervical cord and EMG electrodes were implanted into forelimb muscles. In two cohorts (LIPUS and LIPUS + Buspirone) rats were tested pre-, with- and post-ultrasound stimulation. In LIPUS group rats, fore-limb reaching and grasping success rates first increased and then dropped after 3 weeks while for combination of drug and LIPUS stimulation the score continued to increase. Furthermore, LIPUS stimulation alone did not result in any significant improvement of grip strength compared to the control and combined groups. The findings of this study indicated the potential of LIPUS in SCI recovery and offer a future research direction of a new neuromodulation method.

## 1. Introduction

Spinal cord injury (SCI) is a devastating neuronal dysfunction that affects millions of people each year with significant deficits of motor, sen-sory and autonomic function. According to the World Health Organization (WHO), each year 250,000-500,000 people become newly injured with SCI [1]. The National Spinal Cord Injury Statistical Center (NSCISC), recently reported that incomplete tetraplegia is the most frequent type of SCI [2]. Among the different disabilities in tetraplegic patients, regaining arm and hand function is the highest priority because it could improve their daily quality of life [3]. SCI recovery via neuromodulation that includes electrical [4,5] or pharmacological approaches has recently gained significant popularity [6]. In the rat model, electrical stimulation along with serotonergic agonists have been used to transform non-functional spinal circuits into functional state after loss of supraspinal connections to and from the brain [7-9]. The findings provide an important basis for human studies. In chronic paraplegic and tetraplegic patients, the efficacy of electrical and pharmacological neuromodulation have also been shown to recover upper and lower limb volitional control function [6,10-12]. A better outcome has been recorded with combinatory approaches compared to a single neuromodulator approach alone [13]. However, the most successful electrical stimulation method requires implantation of a stimulator and stimulation electrodes into the patient’s spinal cord.

In recent years significant research has been conducted to elucidate the therapeutic effects of ultrasound stimulation in several neurological disorders such as stroke, Parkinson’s disease and pain management [14]. Because of the non-invasive nature of ultrasound stimulation as well as safety and efficacy considerations, the research interest in this area is in-creasing rapidly [15,16]. The exact mechanism of ultrasound neuromodulation is not known yet. It is assumed that the acoustic force induced by ultrasound generates its biophysical effects on tissues and cells. Therapeutic non-invasive LIPUS stimulation can modulate the neuronal function following brain and peripheral nerve injury because of its high spatial resolution and deep penetration [15]. As a non-invasive method, LI-PUS stimulation is a rapidly growing field for neuromodulation to treat several neurological disorders [17,18] in addition to its conventional ap-plications for bone fracture healing [19].

To the best of our knowledge, therapeutic ultrasound stimulation has not yet been explored in relation to motor recovery in chronic SCI [20,21]. Current study was designed to explore the effects of LIPUS, alone or in combination with a standard pharmacological neuromodulation, Buspirone in rats with C4 cervical cord injury. Forelimb reaching and grasping function, grip strength and distal muscles responses were examined in LIPUS- and LIPUS+Buspirone-treated rats. Similar to electrical neuromodulation, our hypothesis was that LIPUS stimulation can significantly restore or improve forelimb motor functions in rats with cervical cord injured.

## 2. Materials and Methods

### 2.1 Animals

Thirty female Sprague-Dawley rats (3-4 weeks old, weight 230 ± 30 grams b.w.) were used for this study. Because of the low feed conversion ratio and ease of handling, female rats were used in this study. In a poly-propylene cage, the rats were housed with aspen-shaven flooring. The temperature (24°C) and humidity (40%) were strictly monitored in the Centralised Animal Facilities at the Hong Kong Polytechnic University. All experimental procedures were conducted under the guidelines and approval of the Animal Subjects Ethics Sub-committee (ASESC).

### 2.2 Forelimb reaching and grasping

To accustomize the rats with the forelimb reaching and grasping task, during the first 1-2 weeks the rats were handled to familiarize them with a special Plexiglas chamber **(Fig. 1A)** (40 cm × 25 cm × 30 cm; with a 1-2 cm wide opening) for grasping food pellets from a pit [22]. At the same time, the rats were also familiarized with special 45 mg dustless food pellets (Bio-serv®, USA) to grasp and eat by the preferred forepaw. After two weeks of familiarization, the rats were trained to grasp the pel-lets from the pit in a consecutive manner. Food restrictions were provided before starting the training. To master the reaching and grasping task, the pellets were given in the food pit on a pit platform in front of the box slit to ensure that the rats approached the opening in a consistent manner. Ten pellets as a warm-up and 20 pellets per task were usually used to evaluate the reaching behavior. Quantitative assessments of the rats’ skill in the reaching task were performed for six weeks as described previously [22]. The best 26 rats were included for surgery. Following surgery, the rats were subjected to 6 weeks of the motor behavioral task **(Fig. 1B)**.

**Figure 1.**
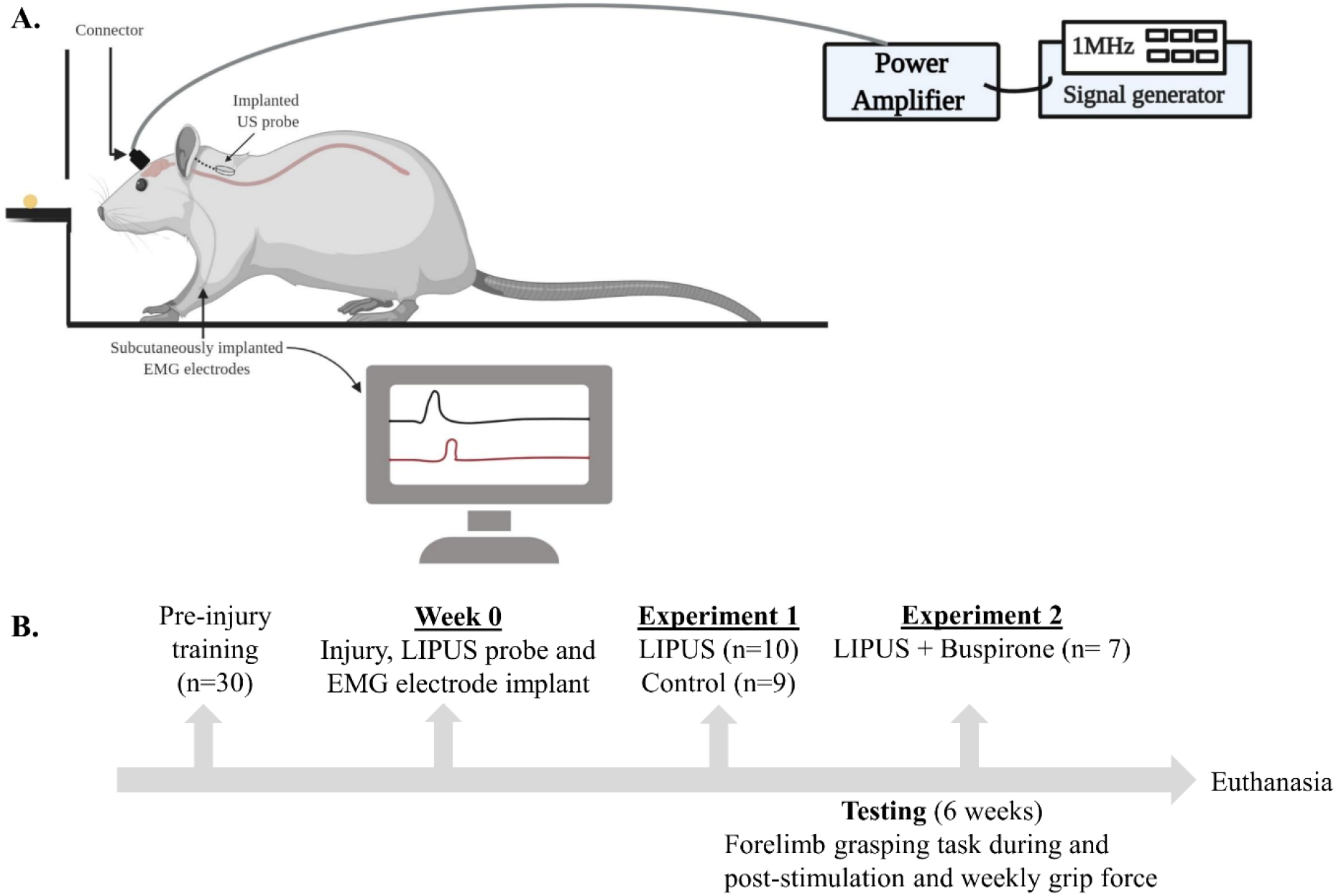
Experimental setup for in vivo LIPUS stimulation in cervical cord in-jured rats. (A) A coaxial cable was connected to a 50 W RF power amplifier to deliver the current to the LIPUS probe that was connected to a SMC connector on the skull (the figure created with biorender.com). (B) Experimental design: Rats (n=30), were trained for 6 weeks to reach and grasp food pellets from a pit platform. After successful training, in the best 26 rats injuries were inflicted at the C4 level and EMG electrodes were implanted in the extensor and flexor digitorum muscles. After recovering from injury, in the first experiment (LI-PUS, n=10 and control, n=9) forelimb reaching and grasping success rates were recorded during and post-stimulation. Each week the maximum grip strength was also recorded to measure the status of quantitative strength of the flexor muscle. In the second experiment additional 7 rats were added to design three groups (LIPUS, LIPUS+Buspirone-treated rats and control). Forelimb reaching and grasping score and the grip forces were recorded and compared to the con-trol group rats.

### 2.3 Grip strength test

The rats were tested and acclimatized with a custom-made grip strength meter as described before [23]. A metal grid was connected to a force sensor and the values were recorded on a computer. By pulling the tail along the horizontal axis the maximum value was recorded and calibrated. At post-injury condition, each week after the forelimb reaching and grasping task the grip strength was recorded. The rats were held in such a way that they could grasp a grid that was connected to a force sensor. The force values for each week were averaged and the maximum strength from pre-injury to week-6 was calculated. The values of each week were normalized from each rat as described before [23][30].

### 2.4 Preparation of LIPUS stimulation probe for implantation

To deliver therapeutic ultrasound in the rats, custom-made implantable LIPUS probes were used in this study. Piezoelectric discs (PZT-8, Beijing Quanxin Ultrasonic Co. Ltd, China) were utilized as the LIPUS probe. To transmit electric power to the piezoelectric disc, two (5-cm each) multi-stranded Teflon-coated stainless-steel wires (AS631, Conner wire Inc., USA) were connected (**Fig. 2A**). The other ends of the wires were connect-ed to the pins of a head connector (SMC) to mount on the head of each rat. The piezoelectric element was then coated with a biocompatible material as described previously [24].

**Figure 2.**
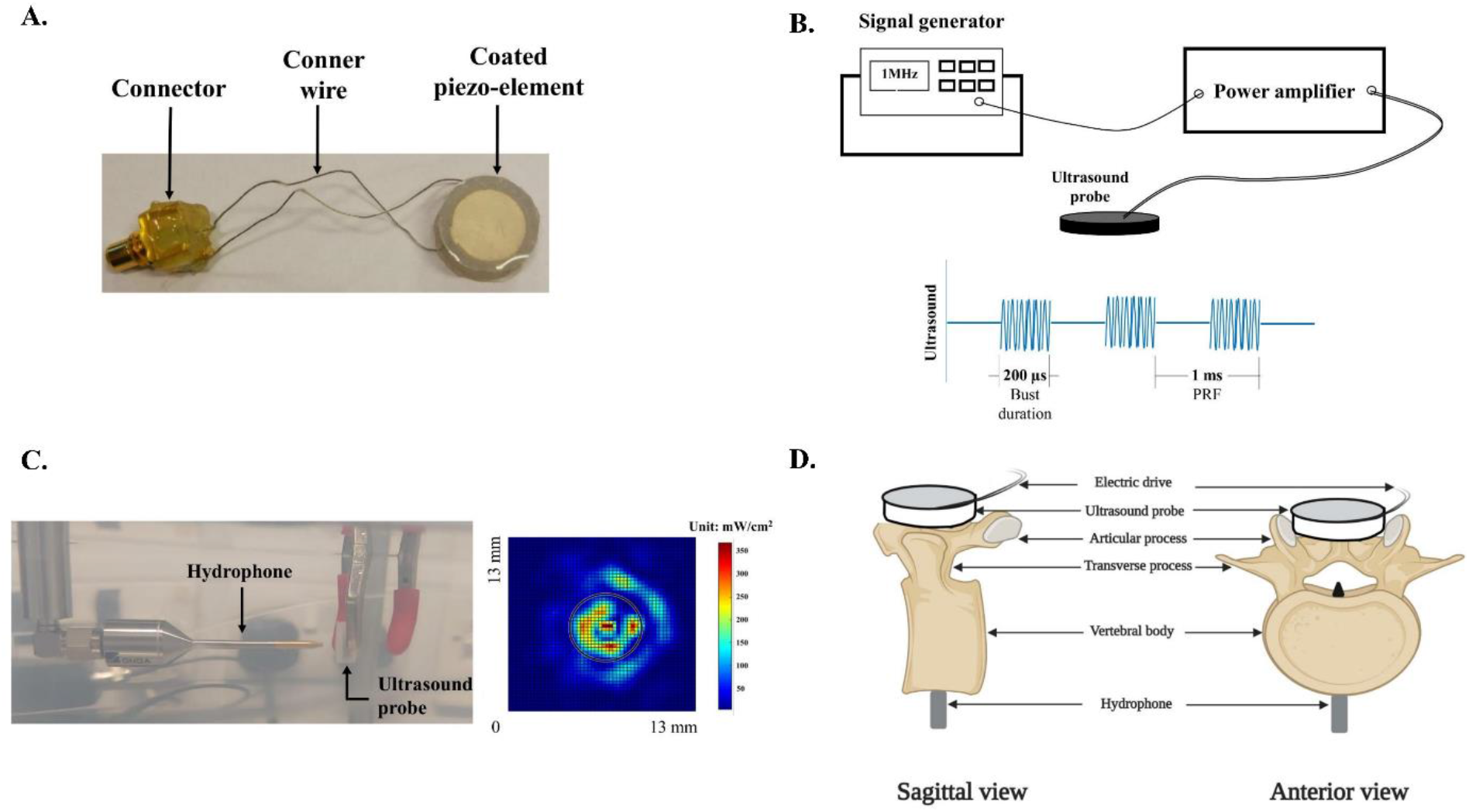
LIPUS stimulation probe and parameters. (**A**) A LIPUS stimulation probe: a piezoelectric element was coated with silicone and connected to a connector via Teflon-coated wires; (**B**) LIPUS stimulation parameter: an acoustic frequency of 1 MHz, 20% duty cycle and 1 ms pulse repetition frequency (PRF) were used to generate the LIPUS. (**C**) Ultrasound intensity was measured in a water tank by placing a hydrophone at the front side of the probe. The in-tensity value was found to be 67.35 mW/cm^2^ (ISATA) (**D**) Ex-vivo setup to measure the ultrasound intensity inside the vertebral canal.

### 2.5 Cervical cord injury

To induce a significant forelimb motor deficit, an incomplete SCI was carried out in 26 rats. The rats were first anesthetized with isoflurane gas (5%) and flow was maintained (1.5-2%) via a facemask throughout the surgery. The rats’ body temperature was maintained at 37 ºC using a heating pad (Thermostar Homeothermic Blanket, RWD^®^, China). Before the surgery, an analgesic Buprenorphine HCL (Buprenex^®^, 0.5 mg/kg, S.C.) was administered. The surgical sites (head and cervical region) were carefully shaved and disinfected by povidone-iodine (Betadine^®^, Mundi-pharma, Switzerland) followed by 70% ethanol. To carry out an incomplete SCI a longitudinal midline incision was made dorsal to the cervical spinal column [25]. Laminectomy was performed by removing fascia and reflecting underlying spinal muscles over the C2-C6 vertebra to isolate the spinous process with the rongeurs. At the C4 level, the spinal cord was exposed by using a bone nibbler. To produce an incomplete injury at the C4 level, the dorsal funiculus was crushed by inserting the tips of fine sharp forceps (2 mm wide and 2 mm deep) as described previously [26]. A hemostat was inserted by placing a small piece of cotton on the incision.

### 2.6 Implanting LIPUS probe and EMG electrode

To anchor the connector of the LIPUS probe on the skull, the rats were placed on a stereotaxic frame. A skin incision was made along the midline of the skull. Fascia and muscles over the skull were reflected laterally. The skull was dried and drilled to put stainless-steel screws into the bone. A SMC connector (RS components^®^, Taiwan) was then placed between the screws. Dental cement was used to fix the connector. An ultrasound probe was then placed at the C4 level of the spinal cord where the dorsal funiculus was crushed and fixed by suturing the wires with the adjunct muscles. The muscles and connective tissue over the cervical region were sutured by using 4.0 Vicryl (ETHICON^®^, NJ, USA) and the skin was closed with a continuous 4.0 Ethilon suture (ETHICON^®^, NJ, USA).

To record the forelimb muscle activities during the reaching and grasping task, EMG electrodes were implanted in the distal forelimb muscles. The objective of implanting the electrodes was to record the extensor and flexor activity before and after the LIPUS stimulation. From each group, two rats were selected to implant the EMG electrodes at the preferred forelimb extensor and flexor digitorum muscles as described previously [23,25]. The head plug of the EMG connector was placed on the skull after retracting the skin and connective tissue. To attach the electrodes to the paw muscles, Teflon-coated stainless-steel wires (AS631, Cooner Wire Inc., USA) were connected to the incised forelimb distal flex-or and extensor digitorum muscle area. Blunt forceps were used to pass the electrodes subcutaneously under the muscle belly. After attaching the electrodes, sharp forceps were used to retract the fascia and locate the muscle belly. To fix the electrodes a 27-gauze needle was inserted into the muscle belly and Teflon-coated wires were inserted into the muscles. A part (∼0.5 mm) of the Teflon from the wires was removed to make EMG electrodes. The electrodes were then anchored tightly at both ends by using 4.0 Ethilon sutures. To confirm the position of the electrodes, electrical stimulation was delivered through the connector to observe muscle contraction. After confirmation, the wires were then coiled subcutaneously to relieve the stress and dental cement was applied to affix the connector.

### 2.7 LIPUS stimulation

For LIPUS stimulation, pulsed ultrasound with a frequency of 1 MHz, duty cycle of 20%, and pulse repetition frequency of 1 kHz was used (**Fig. 2B**) as described in a previous study [27]. A coaxial cable was used to deliver the current from a 50W RF power amplifier to the ultra-sound probe. Before placing the ultrasound probe on the cervical region, the acoustic intensity was measured by using a needle hydrophone (HNP-1000, Onda Corporation, USA). The probe was placed inside a water tank and the hydrophone was placed rostrally to the probe at a 4-mm distance (**Fig. 2C**). To supply the voltage to the ultrasound probe, a coaxial cable was connected to the connector and the intensity was recorded by the hydrophone. To measure the intensity of pulsed ultrasound inside the vertebral canal, an ex-vivo experiment was conducted. First, a vertebra was collected from one rat to measure the intensity inside the vertebral canal (**Fig. 2D**). The vertebral body was drilled by an electric micro driller and the needle of the hydrophone was inserted within the body of the vertebra to measure the intensity of the ultrasound. After 5 minutes of therapeutic stimulation, the behavior task was conducted by using the parameters mentioned above. LIPUS stimulation was delivered for 10 minutes during the reaching and grasping task.

### 2.8 Drug treatment

To determine the combined effect of LIPUS and drugs, in a separate study four rats were administered (i/p) a dose of 1.5 mg/kg b.w. of Buspirone (Tocris®, UK) [23] daily for 6 weeks. Forelimb reaching and grasping success rates and grip strength were recorded during and post-LIPUS stimulation after 30 minutes of Buspirone administration.

### 2.9 Data analysis and statistics

Reaching and grasping success rates were calculated as described before [25,28,29]. A quantitative assessment was performed for each rat. To calculate the success rate, pre-injury 20 pellets were given and post-injury 30 pellets were given (during-stimulation, 20 and post-stimulation, 10). Pre- and post-injury success rates were calculated to find out the difference. A two-tailed paired t-test was used to determine the difference in success rates of reaching and grasping between pre- and post-injury [25]. A two-way analysis of variance (ANOVA) with Tukey post-hoc test was used to determine the difference among the success rates for during- and post-ultrasound therapeutic stimulation. The maximum grip strength from two groups was also analyzed by using a two-way ANOVA with Tukey post-hoc test from week-1 to week-6. For EMG analysis, video footage of reaching and grasping (successful and unsuccessful) attempts was examined frame by frame via a media player. The EMG signals were bandpass filtered at 10-1000 Hz and amplified 1000 times using an ana-log amplifier (Model 1700 Differential AC Amplifier, AM Systems, USA). A data acquisition system (Power1401-3A, Cambridge Electronics Design Ltd., UK) was used to digitize the EMG signals. A software (Signal, Cam-bridge Electronics Design Ltd., UK) was used to visualize the EMG signals and synchronize them with the video during the reaching and grasping task. The EMG data of extensor and flexor muscles were normalized as described before [23,25]. The difference between groups was considered as significant if *p*<0.05. Statistical analysis was performed using Prism (GraphPad Prism Software, version 8.4.2, USA) and MATLAB (Math Works Inc., Natick, USA).

## 3. Results

### 3.1 Therapeutic intensity of the LIPUS probe

In water at a 4-mm distance, the average ultrasound intensity of the LIPUS probe was found to be 67.35 mW/cm^2^ (I_SATA_) (**Fig. 2C**). From the ex-vivo experiment inside the spinal canal, the average intensity of the ultra-sound signal reaching the spinal cord area was found to be 32 mW/cm^2^. For the LIPUS treated rats, the same intensity of LIPUS probe was used to find out the therapeutic effect. The stimulation was provided for about 10 minutes [25] during the reaching and grasping task and success rates were recorded.

### 3.2 LIPUS stimulation facilitates the forelimb reaching and grasping function

Following injury, the rats lost their grasping function (pre-injury vs. post-injury day-8; 73.95 ± 1.301 vs. 3.68 ± 2.158, \*\*\**p*<0.001, paired t-test). After one week of recovery from injury, the LIPUS-treated rats were tested for forelimb reaching and grasping task and success rates were compared with the control group. A total of 19 animals (LIPUS, n=10 and control, n=9) were included to evaluate the forelimb reaching and grasping function as well as the grip strength.

At week-2 and week-3 during stimulation, improvement of reaching scores was found compared to the control group. The post-stimulation success rate was also found to be higher than the control group scores at week-2 and week-3. The success rates in the LIPUS group rats were found to be higher compared to those of the control group rats (no stimulation but implanted with the same stimulation probes) from week 1 to 6. The LIPUS group rats had significant improvement at week-2 (30.00 ± 6.28 vs. 5.92 ± 2.20; p =0.0079), (30.16 ± 7.31 vs. 5.92 ± 2.20; p =0.0074) and week-3 (41.50 ± 3.11 vs. 15.92 ± 3.92; p =0.0044), (37.167 ± 4.54; p =0.0223) com-pared to the control group rats (**Fig. 3A**). However, from week-4 onward no significant improvement was found. At week-6 little more improvement was observed during (44.16 ± 4.98) and post (38.66 ± 6.15) stimulation, compared to the control group (26.30 ± 5.76). However, the score at week-6 was not significant like week-2 and week-3 scores.

**Figure 3.**
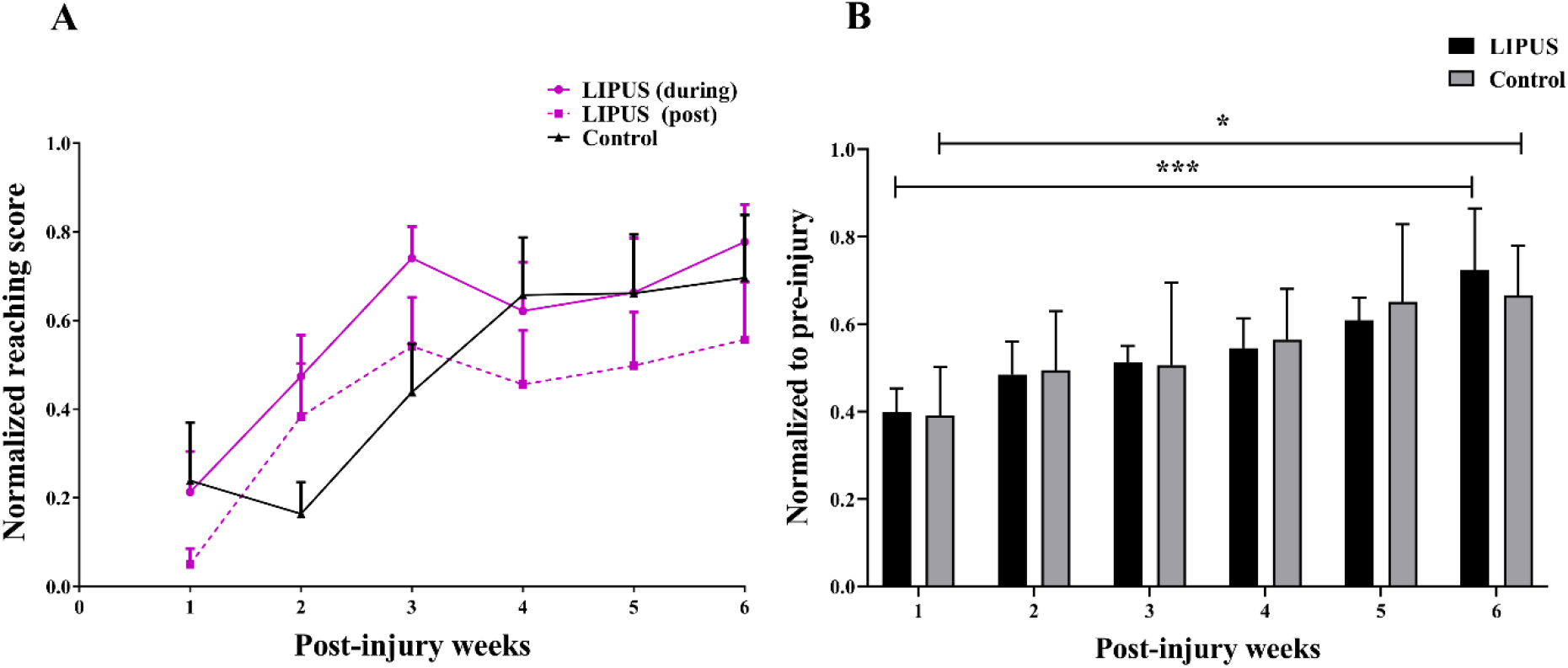
Behavioral success rate; (**A**) Normalized reaching score (LIPUS, n=10 and control, n=9) of the reaching and grasping task in LIPUS and control group rats. The solid pink line indicates the scores when the LIPUS rats were receiv-ing the stimulation (20 trials/rat/test session) and the dotted pink line repre-sents the scores immediately after the stimulation (10 trials/rat/test session). The black line indicates the success rates of the control group rats that did not receive ultrasound stimulation (20 trials/rat/test session). (**B**) Normalized grip strength (LIPUS group n=6 rats and control group, n=5). Data are presented as mean and SEM (LIPUS group, n=6; and Control group, n=5). Grip strength values were normalized to the maximum values. Significant improvements were found in the LIPUS group rats (\*\*\**p* < 0.001, two-way ANOVA, Tukey post-hoc test) and in the control group at week-6 (\*\**p* < 0.01, two-way ANOVA, Tukey post-hoc test) compared to week-1 post-injury.

### 3.3 LIPUS improves forelimb grip strength

Following the skilled reaching and grasping test, the grip strength of 11 rats (LIPUS group, n=6 rats, and Control group, n=5 rats) was deter-mined by using a custom-made grip strength meter as described before [30]. Significant muscle strength improvements were found at week-6 compared to week-1 post-injury in the LIPUS group (**Fig. 3B**). However, in the control group, slightly significant improvements were found at week-6. In addition, no significant differences were observed between the two groups at any week.

### 3.4 Effects of LIPUS on EMG synergy

During reaching and grasping EMGs were recorded from the extensor digitorum and flexor digitorum muscles at 1-to 3-week post-injury and the control group rats were recorded at 1- and 6 weeks post-injury. The raw EMG signals from the extensor and flexor digitorum muscles are presented in **Fig. 4**. EMG signals of the extensor digitorum and flexor digitorum muscle values were normalized to calculate the AUC. Values were compared to the pre-stimulation and post-stimulation at week-1, -2 and -3 (**Fig. 5**).

**Figure 4.**
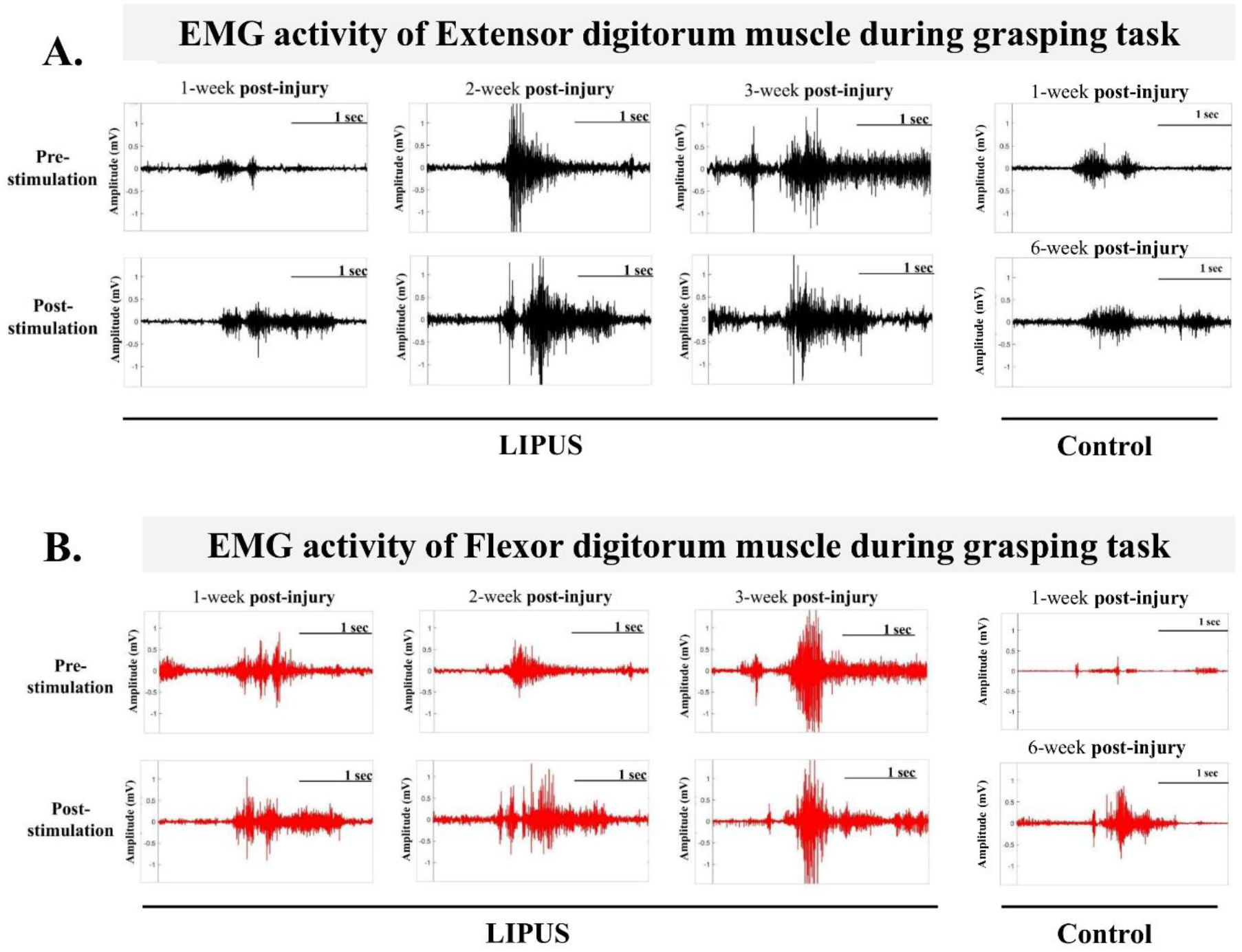
Raw EMG signals of (A) extensor digitorum (black) and (B) flexor digitorum (red) muscles during the forelimb reaching and grasping task of the LIPUS group rats at three different post-injury weeks. and control group rats at week-1 and week-6 post-injury. One representative rat EMG is presented in the figure.

**Figure 5.**
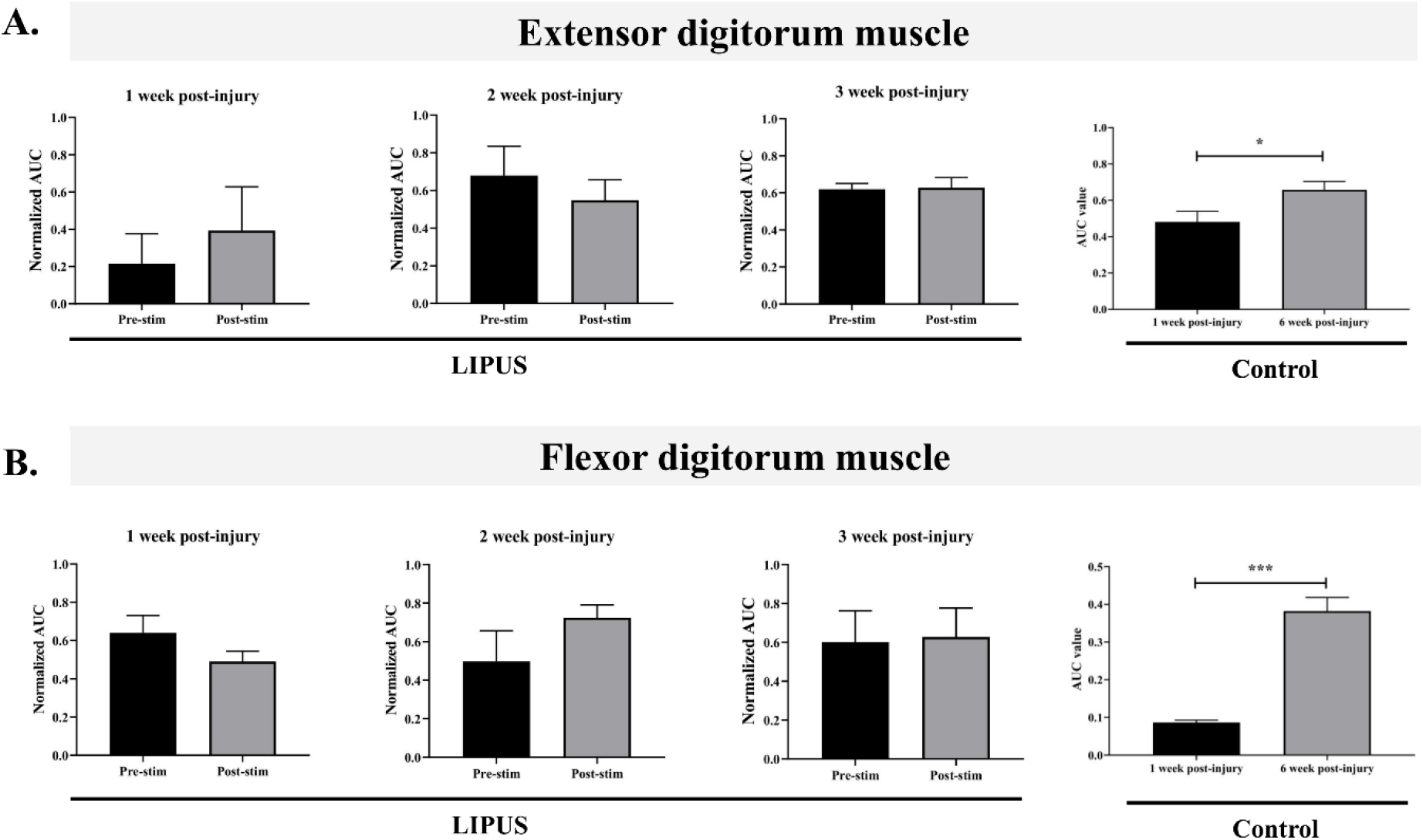
Normalized area-under-the-curve (AUC) of EMG signals from (**A**) ex-tensor digitorum and (**B**) flexor digitorum muscles of LIPUS group rats (n=2) and control group rats (n=1) during the single pellet reaching task (10 trials/rat). AUC value of LIPUS group rats were recorded at week-1, week-2 and week-3 post-injury and the AUC value of control group rats was recorded at week-1 and week-6 post-injury. Significant differences were observed in extensor muscle (\**p* < 0.05, unpaired two-tailed t-test) and flexor muscle (\*\*\**p* < 0.001, unpaired two-tailed t-test). Data are presented as mean and SEM.

In contrast, for the flexor muscle at week-1 the values decreased after stimulation and increased at week-2. Like the extensor muscle, in the flexor muscle no difference was found at week-3. This indicates that 3-week post-injury stimulation could not facilitate the reaching and grasping task unlike week-1 and week-2. However, no statistically significant difference was found at 3 different weeks. From **Fig. 5** it is observed that the control group rat muscle amplitude increased without any stimulation after injury similar to as found before. The raw EMG signals are presented in **Fig. 5**. In the control group at week-6 post-injury the AUC value increased from week-1 in both extensor (0.48 ± 0.05 to 0.65 ± 0.04, \**p*<0.05, unpaired two-tailed t-test) and flexor (0.09 ± 0.006 to 0.38 ± 0.03, \*\*\**p*<0.001, unpaired two-tailed t-test) muscles (**Fig. 5**).

### 3.5 Combined therapy further improves forelimb fine motor function and grip strength

The success rate in the LIPUS stimulation group and combined group were found to be higher compared to the control group’s reaching and grasping success from week-1 to week-6 post-injury treatment. A significant improvement was found at week-3 during ultrasound stimulation (39.167 ± 3.96 vs. 11.667 ± 5.27; p =0.0079) (**Fig. 6A**). However, after week-3 the success rate dropped in the LIPUS stimulation group and did not exhibit any significant improvement until week-6. Conversely, in the combined group, rats achieved significantly improved forelimb function-al recovery in a consistent manner compared to the LIPUS stimulation groups, at week-5 and week-6 (48.75 ± 8.98 vs. 12.50 ± 5.73; p =0.0097 and 50.00 ± 10.80 vs. 15.00 ± 6.83; p =0.0139) during stimulation compared to the control group. Post-stimulation, the improvement was also found significantly higher at week-5 and week-6 compared to the control group (50.00 ± 10.00 vs. 12.50 ± 5.73; p =0.0067 and 52.50 ± 11.08 vs. 15.00 ± 6.83; p =0.0067).

**Figure 6.**
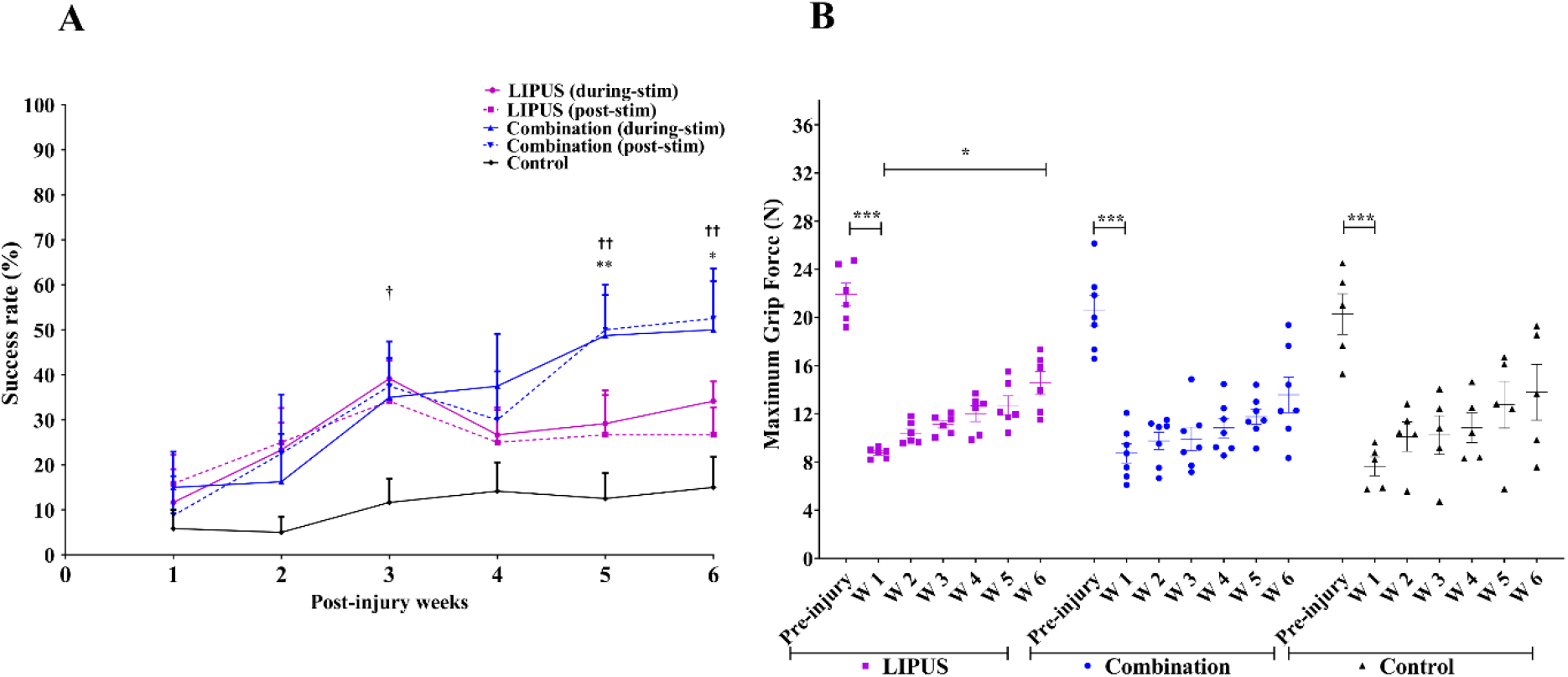
(**A**) Average success rate (mean ±SEM; LIPUS, n=6, combined, n=4 and control, n=5) of reaching and grasping tasks of three groups of rats. The solid pink line indicates the success rates of the ultrasound group rats when receiving the ultrasound stimulation and the dotted line is the success rates immediately after stimulation. The solid blue line indicates the success rates of the combination group rats when receiving the ultrasound stimulation and the dotted line is the success rate immediately after stimulation. The black line indicates the success rate of the control group of rats that did not receive any stimulation or drug treatment. At week-3 during ultrasound stimulation, significant (†p < 0.05, two-way ANOVA, Tukey post-hoc test) improvement of reaching score was found in the LIPUS stimulation group compared to the control group. During stimulation the success rate of the combined group was found to be significantly higher compared to the control group at week-5 and week-6 (\*\**p* < 0.01 and \**p* < 0.05 two-way ANOVA, Tukey post-hoc test). How-ever, at week-5 and week-6, the poststimulation success rate was found to be significant (^††^*p* <0.01, two-way ANOVA, Tukey post-hoc test) compared to the control group rats. (**B**) Maximum grip strength of three different groups (mean ±SEM; LIPUS, n=6, combined, n=7 and control, n=5). At week-1 post-injury, the grip strength dropped significantly compared to the pre-injured condition in three groups of rats (LIPUS group, 21.93 ± 0.93 vs. 8.75 ± 0.17; combined group, 20.55 ± 1.24 vs. 8.22 ± 0.79 and control group 20.28 ± 1.68 vs. 7.63 ± 0.78; \*\*\**p* < 0.001, two-way ANOVA, Tukey post-hoc test). At week-6 the grip strength of LIPUS group rats improved (\**p* < 0.05 two-way ANOVA, Tukey post-hoc test) compared to that at week-1 post-injury.

The maximum grip strength of LIPUS group rats was found to be 8.75 N (week-1) to 14.56 N (week-6). For combined group rats, the value was from 8.72 N (week-1) to 13.58 N (week-6). In the control group rats there was a similar increasing trend, from 7.63 N (week-1) to 13.79 N (week-6) (**Fig. 6B**).

### 3.6 Combination therapy altered the muscle coordination in distal muscle

EMG activity was observed from one representative rat at week-1 and week-5 post-injury (**Fig. 7**). To calculate the normalized AUC of EMG signals of the extensor and flexor digitorum muscles, the values were normalized. From the LIPUS and combined groups, the normalized values were compared to the pre- and post-stimulation with the control group at week-1 and week-5 post-injury (**Fig. 8**). At week-1 post-injury, the normalized value of the extensor muscle decreased after simulation in the combined and LIPUS groups. Moreover, a significant decrease of muscle activity was found at week-5 post-injury in the extensor muscle compared to the pre-stimulation (0.82 ± 0.09 vs. 0.15 ± 0.05; p= 0.0025) (**Fig. 8A**). A similar response was found in the flexor muscle at week-1 in the extensor muscle. However, at week-5 post-injury the flexor muscle values in-creased in the LIPUS group and decreased in the combined group rats after stimulation (**Fig. 8B**). At week-1 and week-5 post-injury EMG signals were recorded from three representative rats.

**Figure 7.**
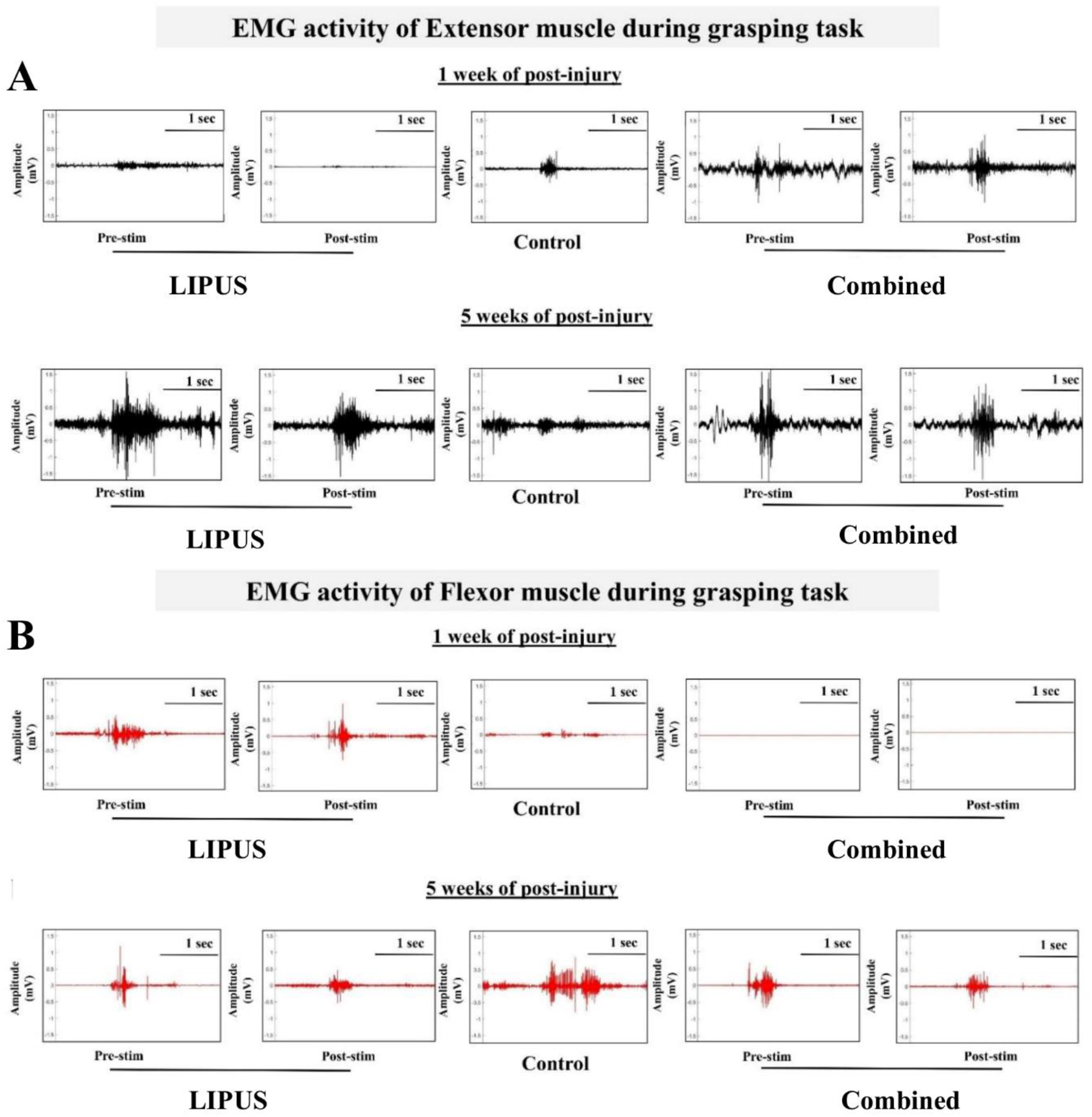
Raw EMG signals from three representative rats (**A**) Extensor digitorum and (**B**) Flexor digitorum muscle at week-1 post-injury and week-5 post-injury.

**Figure 8.**
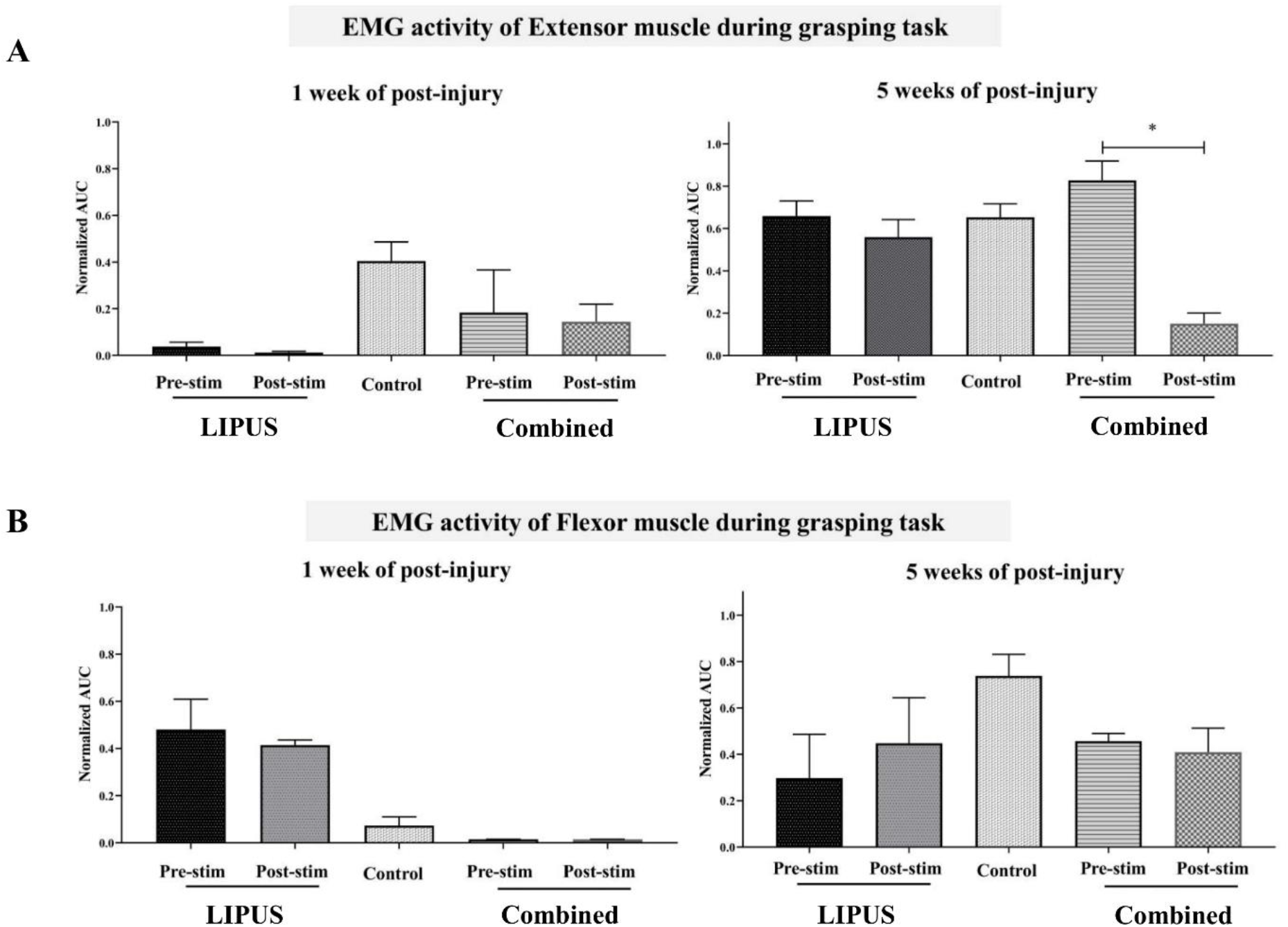
Normalized area-under-the-curve (AUC) of EMG signals from (**A**) ex-tensor digitorum and (**B**) flexor digitorum muscles of three group of rats dur-ing the single pellet reaching task (n=10 trials). A significant difference was found at week-5 post-injury in the combined group of extensor muscle (\**p* < 0.05 two-way ANOVA, Tukey post-hoc test). Data are presented as mean and SEM. (LIPUS group, n=2; Combined group; n=1 and Control group, n=1).

## 4. Discussion

In the present study, the effect of LIPUS was evaluated via a forelimb grasping task and grip strength in chronic incomplete spinal cord injured rats. Moreover, two different antagonistic muscle responses were also evaluated. An incomplete cervical injury at the C4 level exhibited a significant deficit in motor function because of the disruption of the motor neuronal network. The synergistic effect of LIPUS and serotonergic drug indicates a significant role in modulating neural networks after cervical cord injury and improving functions.

LIPUS is one of the choices of neuro-stimulation because of its penetration and modulation capability in the neurons [15]. In our experiment, a skilled motor reaching and grasping task was used to evaluate and compare different conditions: during and post-ultrasound stimulation. The effects of ultrasound stimulation on the success rate were found to be higher during stimulation compared to post-stimulation. The maximum success rate of LIPUS stimulation group rats was found at week-3; how-ever, afterward, the success rates decreased (**Fig. 3A**). At week-3 post-injury, no significant difference was found, although the success rate was higher compared to the control group of rats.

The possible cause is that after 3 weeks astrocytic scarring matures and converts to a fibrotic scar [31] and ultrasound cannot facilitate further recovery behind the chronic scar. Moreover, in a previous study it has al-ready been shown that after three weeks, ultrasound-driven piezoelectric voltage drops because of the acoustic impedance of the growing scar around the ultrasound probe [24]. In our early observation in LIPUS group rats the forelimb reaching and grasping success rate dropped after 3 weeks (**Fig. 3A**). The same events may have happened in our subsequent LIPUS group rats (**Fig. 6A**). In another aspect, during and post-ultrasound stimulation, no significant differences were found throughout the period. The possible mechanism in the LIPUS group rats is the early neuromodulatory effect of ultrasound that modulates the inter-neuronal activity. In combination group the success rate increased in a continuous manner until 8 weeks. The possible mechanism of last 4 weeks of recovery may be mainly coming from the effect of Buspirone [23].

Like other neuromodulation strategies, the stimulation effect may last for several minutes post-stimulation [25]. To answer this critical question, after turning off the ultrasound stimulation, the reaching score was measured immediately. This scoring lasted up to about 5 min post-stimulation. The non-significant difference between the scores during and post-stimulation suggest that the stimulation was neuromodulatory. To get the most from the neuromodulation, even for the combination group rats, ultrasound stimulation was given for 5 min before the reaching task. It is expected that the time is enough to neuromodulate the cervical cord and improve forelimb function. The combination group rats exhibited consistent recovery in the forelimb reaching and grasping task.

Forelimb grip strength measurement is a useful tool for measuring the recovery for incomplete injury [32]. Like the forelimb reaching and grasping task, the task is not skilled. The distal flexor muscle is responsible for the grasping and gripping ability of rats that are controlled by corticospinal tract [22]. Moreover, the gripping ability is also determined by some part of the reticulospinal tract that descends in the medial part of the ventral column. From our experiment, it is also noticeable that after partial injury the control group of rats recovered their gripping ability. The finding is similar to the result published by Anderson and co-workers [32]. Following injury, the forelimb muscles lose their grasping function because of no supraspinal input to the flexor muscle. However, other forelimb muscle function is not affected significantly after the injury. Hence, the grip strength result is not consistent with the forelimb reaching and grasping task. Moreover, in our experiment, normalized grip strength data showed that the strength tends to increase in the LIPUS group, and a significant improvement was found at week-6 compared to week-1 (**Fig. 3B**).

The EMG signals were found reliable in only few rats throughout the study period and were included for the analysis. In the future study, more animals need to be examined on their electrophysiological changes in response to LIPUS treatment. In our study, the EMG data show that at week-6 in the control group following injury the muscle activation increased compared to the pre-injury condition that was previously described [25]. The hyperexcitation condition of the neural network may cause more energy facilitation in the distal forelimb muscles com-pared to immediately after injury. Moreover, in the LIPUS group the antagonistic activity of the two different extensor and flexor muscles was found at week-1 and week-2 post-injury. Flexor muscle activity increased after stimulation at week-2 indicated by the increase of grasping rate similar to the forelimb reaching and grasping success rate (**Fig. 5**). However, after 2 weeks post-injury, no differences were observed in extensor and flexor muscles before and after stimulation.

Spinal cord neuromodulation via LIPUS is still an unexplored avenue in SCI recovery research. Furthermore, combined therapy of non-invasive neurostimulation by LIPUS and serotonergic neuromodulation is a unique approach. In a combined neuromodulation approach, fore-limb reaching and grasping success rate was found to be higher com-pared to the LIPUS group, which indicates that drug-based neuromodulation has a significant role to play in neural recovery (**Fig. 6A**). Although no significant difference was found at week-6 in grip strength (**Fig. 6B**), in LIPUS groups a significant improvement was found at week-6 compared to week-1 (**Fig. 6B**). A significant improvement was also found at week-5 post-injury in the extensor digitorum muscle post-stimulation. However, flexor muscle did not exhibit any response.

The present study, however, has a few limitations. The overall intensity of all custom-made ultrasound probe were in the range of 60-70mW/cm^2^. But the LIPUS intensity of the ultrasound probes were not controlled and maybe variable in different implants. With a reliable in vivo testing of LIPUS intensity, the therapeutic results will be more accurate. In addition, the thermal effect of ultrasound stimulation was not considered negligible in this study, which could have some potential effects on the recovery.

Furthermore, histopathological examination of the spinal cord after LIPUS stimulation was not done in the current study. Histopathological data can further provide important information on the neuropathological changes of the spinal cord circuits after the LIPUS treatment. In a recent study, it has been shown that low-intensity ultrasound stimulation could induce the hindlimb recovery following T10 contusion SCI in a rat model, where MR imaging and histological findings showed significant reduction of the edema and inflammatory responses (iNOS and TNF-α) with a decreased macrophage marker [33]. Similarly, in mouse traumatic brain injury model transcranial LIPUS stimulation significantly attenuates edema and contused volumes [34]. In peripheral nerve injury, therapeutic ultrasound suppresses the expression of inflammatory cytokines, nerve growth inhibitor [35], promote myelination and axonal regeneration [36] following in rat model. Above findings from other studies suggest that LIPUS has neuropathological effects and is a potential therapeutic approach for neural regeneration. Further studies are needed to investigate the histopathological changes in the spinal cord after LIPUS therapy.

## 5. Conclusions

The findings of this research include possible functional recovery by ultrasound spinal cord stimulation and a definite recovery of forelimb function when the ultrasound stimulation is combined with serotonergic agonist drug-based neuromodulation in incomplete cervical cord injured rats. Finding recoveries after the loss of upper extremity function due to a cervical injury in recent years has drawn more attention because of their high clinical significance. Following skill training, rats used their forelimbs for feeding after successful grasping of food. The reaching and grasping behavior in rats is quite similar to that in humans. This allows the behavioral task to be a powerful tool to translate in clinical conditions. However, the finesse of digit control is less developed in rodents than in non-human primates. The use of non-human primates could be more useful to investigate the behavioral and electrophysiological changes.

